# Benchmarking Computational Integration Methods for Spatial Transcriptomics Data

**DOI:** 10.1101/2021.08.27.457741

**Authors:** Yijun Li, Stefan Stanojevic, Bing He, Zheng Jing, Qianhui Huang, Jian Kang, Lana X. Garmire

## Abstract

The increasing popularity of spatial transcriptomics has allowed researchers to analyze transcriptome data in its tissue sample’s spatial context. Various methods have been developed for detecting SV (spatially variable) genes, with distinct spatial expression patterns. However, the accuracy of using these SV genes in clustering has not been thoroughly studied. On the other hand, in single cell resolution sequencing data without spatial context, clustering analysis is usually done on highly variable (HV) genes. Here we investigate if integrating SV genes and HV genes from spatial transcriptomics data can improve clustering performance beyond using SV genes alone. We examined three methods that detect SV genes, including Giotto, spatialDE, and SPARK, and evaluated six methods that integrate different features measured from the same samples including MOFA+, scVI, Seurat v4, CIMLR, SNF, and the straightforward concatenation approach. We applied these methods on 19 real datasets from three different spatial transcriptomics technologies (merFISH, SeqFISH+, and Visium) as well as 20 simulated datasets of varying spatial expression conditions. Our evaluations show that MOFA+ and simple concatenation have good performances in general, despite the variations among datasets and spatial transcriptomics platforms. This work shows that integrating highly variable and spatially variable genes in the spatial transcriptomics data can improve clustering beyond using spatially variable genes only. It also provides practical guides on the choices of computational methods to accomplish this goal.

## Background

Spatial omics technologies are one of the breakthroughs in science in the last several years [1], [2–4]. Such technologies are able to measure transcriptome information systematically in the tissue space, thus preserving the spatial context of the tissue samples. The addition of spatial information allows researchers to further explore biological architecture and function and reveal more insights with respect to various disease mechanisms [5–7] [8] [9]. Various techniques for sequencing spatially resolved transcriptome data have been developed, including merFISH [10] [11] [12], SeqFISH [13], SeqFISH+ [14], Visium [15] [16,17], etc. Such technologies can be categorized into two general classes: fluorescence in situ hybridization (FISH)-based methods [14] [10], which directly extract transcriptome information at a molecular level and obtain the spatial locations of the cells by processing the FISH images; and array-based methods [16] [15], which attach probes with fixed physical locations to cryosections of tissues to obtain transcriptome information.

Since the function of cells is often related to their location in the tissue and their genetic information, spatial transcriptomics methods have great potential in helping us gain insights into biological mechanisms. Genes with spatially distinct expression patterns, or spatially variable (SV) genes, are a novel type of markers for identifying cell types. Recently several computational methods, such as trendsceek [18], spatialDE [19], SPARK [20], Giotto [21], and MERINGUE [22], have been developed with the function to identify spatially variable (SV) genes. However, conventionally clustering and cell-type identification are based on highly variable (HV) genes, as those done in single cell RNA-Seq datasets which provide no spatial information directly. In spatial transcriptomics data, the SV genes with similar spatial expression patterns are often used to segment cells / spots into spatial domains or clusters [16,19]. Notably, the SV genes and HV genes are often quite distinct sets, despite the possibility of overlapping on some genes. We therefore asked if the integration of conventional HV genes and the novel SV genes (both from spatial transcriptomics data), can help to improve clustering performance of this new type of data, an area unexplored currently.

Towards this goal, we conducted a comprehensive benchmarking study to integrate HV genes and SV genes in spatial transcriptomics data. We adapted six different computational methods for this task, namely scVI, CIMLR, SNF, Seurat v4, MOFA+, and the straightforward concatenation approach. scVI is a neural-network based method originally designed for integrating single-cell transcriptomic data [23]. Seurat v4 is a Weighted Nearest Network method [24] and MOFA+ is a matrix factorization method [25], both of which were used to integrate multi-omics single cell data[26]. We also adopted two additional methods originally developed to integrate bulk multi-omics data: CIMLR (Cancer Integration via Multikernel Learning) model is an integration method using Gaussian kernels [27] and SNF is a network fusion method based on patient-patient similarities [28]. While other methods align multiple types of genomics data from similar (but not the same) cells, such as LIGER [29], MATCHER [30] and Harmony [31], these methods are deemed unsuitable for the question in the study, since the SV genes and HV genes here are obtained from the same cells (or spots).

We obtained SV genes using three different popular spatially variable genes detection methods: Giotto [21], spatialDE [19], and SPARK [20]. We then applied the aforementioned integration methods to combine HV and SV genes on a total of 19 real datasets across three spatial transcriptomics platforms, including the Mouse Somatosensory Cortex dataset by SeqFISH+[14], six datasets by 10X genomics’ Visium platform [16,32–36] and twelve datasets by merFISH methods [10]. We also evaluated the clustering results on 20 simulated datasets. The clustering performance was measured using three primary metrics: AMI (Adjusted Mutual Information), which evaluates the accuracy of clustering in spatial omics data by simply comparing the clustering labels to the ground truth labels, without taking into account the spatial structure of the cells / spots; VN (total Variation Norm), which evaluates accuracy by comparing the clustering labels to the ground truth taking the spatial structure of the cells / spots into consideration; and F-1 score which evaluates the accuracy of downstream differential expression analysis. We also assessed the methods’ performance through secondary practical metrics such as running time and memory use. The benchmarking results are expected to provide practical guidelines for researchers conducting spatial omics clustering analysis, where the SV genes and HV genes present distinct properties of the data.

## Results

### Overview of Benchmark Comparisons

We conducted a comprehensive benchmarking study to evaluate the effect of integrating HV genes and SV genes in spatial transcriptomics data on 19 real datasets and 20 simulated datasets, using 5 different integration approaches. The characteristics of the integration methods used in the study are summarized in **Table 1**. We strictly selected only methods that integrate multi-omics data generated from the same cells and excluded methods that align multiple types of genomics data from similar (but not the same) cells, since the SV genes and HV genes here are obtained from the same cells (or spots). The processing pipeline for benchmarking is shown in **Fig. 1**. It starts with spatial transcriptomics data which has two components: a gene expression matrix, and spatial data which consists of the spatial coordinates of each spot or cell. After data preprocessing and normalization (see **Methods**), we extracted the HV genes using the gene expression matrix and the SV genes using both the gene expression matrix and spatial coordinates data. Since HV genes and SV genes normally overlap partially, here we define HV genes as genes that are uniquely highly variable genes, to avoid double-counting the overlapped genes during integration. We then conducted dimensionality reduction to obtain the same number of extracted features, using either the internal steps within the methods (MOFA+, scVI, Seurat v4 and CIMLR) or using PCA (Principal Component Analysis) for the approach without this step (concatenation). Next, we applied clustering (default Leiden algorithm) for all methods except for CIMLR, which has its own built-in clustering step. For a fair comparison of evaluation metrics, we repeatedly performed the Leiden clustering algorithm, tuning its parameters until we obtained the same number of clusters as those from the ground truth. Finally, the clustering performance was assessed by conventional clustering metric AMI (Adjusted Mutual Information) and the novel spatial clustering metric VN (Variation Norm). AMI simply measures the level of concordance between clustering labels and the ground truth labels; VN calculates the concordance between clustering labels and ground truth labels while taking into account the spatial distribution of the cells / spots, a unique feature of spatial transcriptomics data. We also performed differential expression analysis based on the clustering labels using the F-1 score, to assess the downstream quality of the clustering labels. Since SV genes are often used as features in downstream spatial transcriptomics data analysis, we used the results obtained from using SV genes as the baselines, in comparison to those obtained from integration methods.

**Figure 1.**
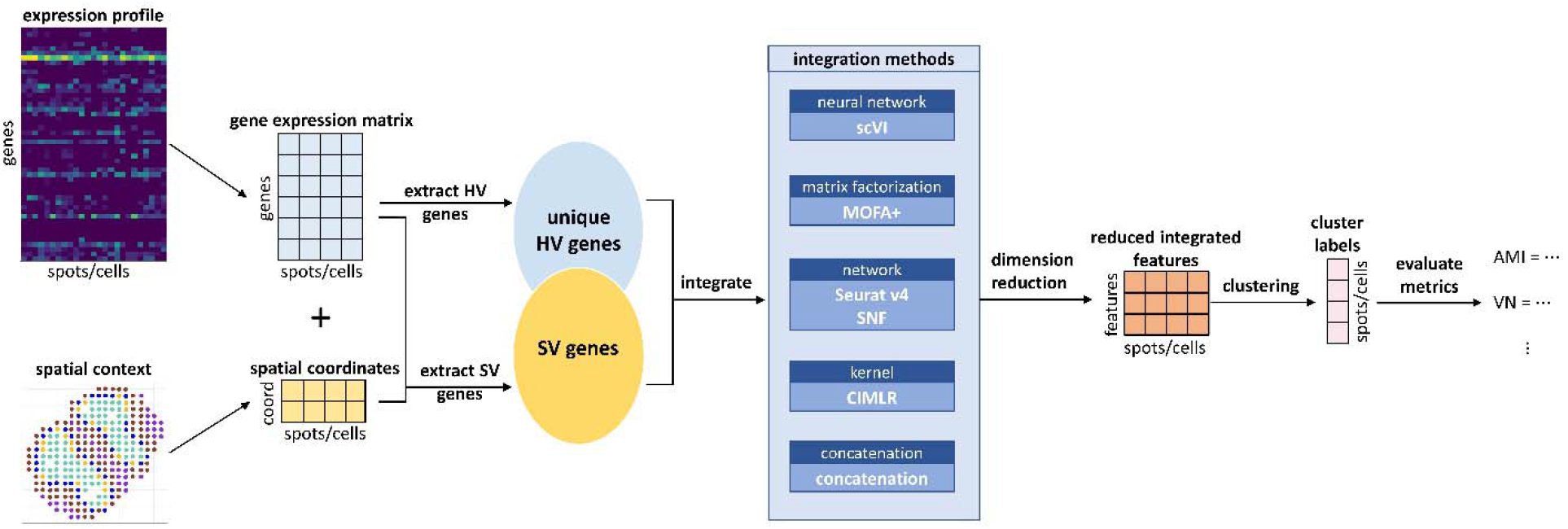
Benchmark workflow. The workflow is composed of three general steps. Step 1: extraction of the HV and SV genes. Step 2: integration and dimension reductions of the HV and SV genes. Step 3: clustering analysis and evaluations based on the integration results.

### Clustering Accuracy on Real Spatial Transcriptomics Datasets

We compared the clustering accuracy performance of paired multi-omics integration methods on 19 datasets with available ground truth information. Our real datasets span over diverse resolutions, tissue types and cell type compositions (**Table 2**). Six datasets are from Visium [15], a non-single-cell resolution technology, or its prototype technology. They include a Mouse Olfactory Bulb dataset [16], a Human Cerebellum dataset [32], a Mouse Kidney dataset [33], and three different Mouse Brain datasets [34–36]. We included the Mouse Somatosensory Cortex dataset from SeqFISH+ [14], a single-cell resolution dataset. We also selected twelve datasets on separate layers of the Mouse Hypothalamus by merFISH [10], another single-cell resolution technology.

To evaluate clustering accuracy, we first compared the methods’ performance using the conventional metric AMI (Adjusted Mutual Information). We compared the clustering results with those using SV genes as the baseline (**Fig. 2a-b, Supplementary Fig. 1a-c**). While there is no single best integration method for all datasets, there is a strong trend across different SV genes detection methods that MOFA+ and the simple concatenation approach yield the best results in general (**Fig. 2a-b, Supplementary Fig. 1a-c**). On average, across different SV genes selection methods, concatenation (p-value = 0.004, 0.049, and 0.002 for Giotto, spatialDE, and SPARK) and MOFA+ (p-value = 0.018, 0.395, and 0.009 for Giotto, spatialDE, and SPARK) have the highest median AMI values across datasets (**Fig. 2a**), significantly better than the baseline. They are followed by scVI, which also have higher AMIs than clustering solely based on SV genes, however the advantage is not statistically significant. Oppositely, SNF (p-value = 7.6e-06, 3.8e-06, and 3.8e-06 for Giotto, spatialDE, and SPARK), CIMLR (p-value = 3.8e-06 for Giotto, spatialDE, and SPARK) and Seurat v4 (p-value = 0.008, 0.001, and 3.4e-04 for Giotto, spatialDE, and SPARK) are significantly worse at integrating SV and HV genes, compared to the baseline of spatial omics clustering using SV genes (**Fig.2a**). In particular, Seurat v4 has a much wider range of AMIs compared to other methods, prompting a closer look at the AMI values across different datasets. Indeed, Seurat v4 shows technology-dependent performance patterns (**Fig. 2b, Supplementary Fig. 1a-c**). It has some of the lowest AMIs in datasets of single cell resolutions, particularly in merFISH datasets; However, it is ranked better in the Visium platform.

**Figure 2.**
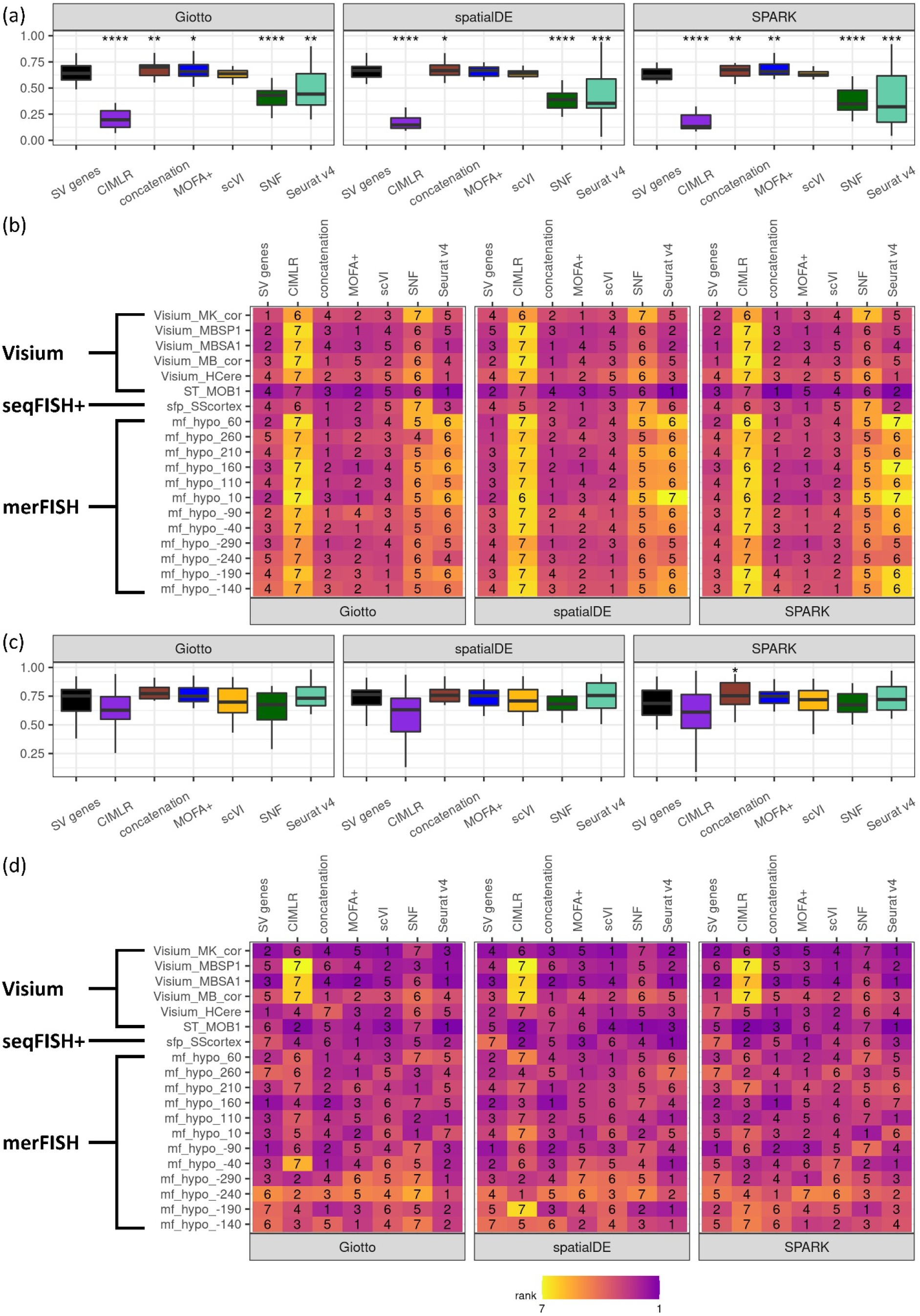
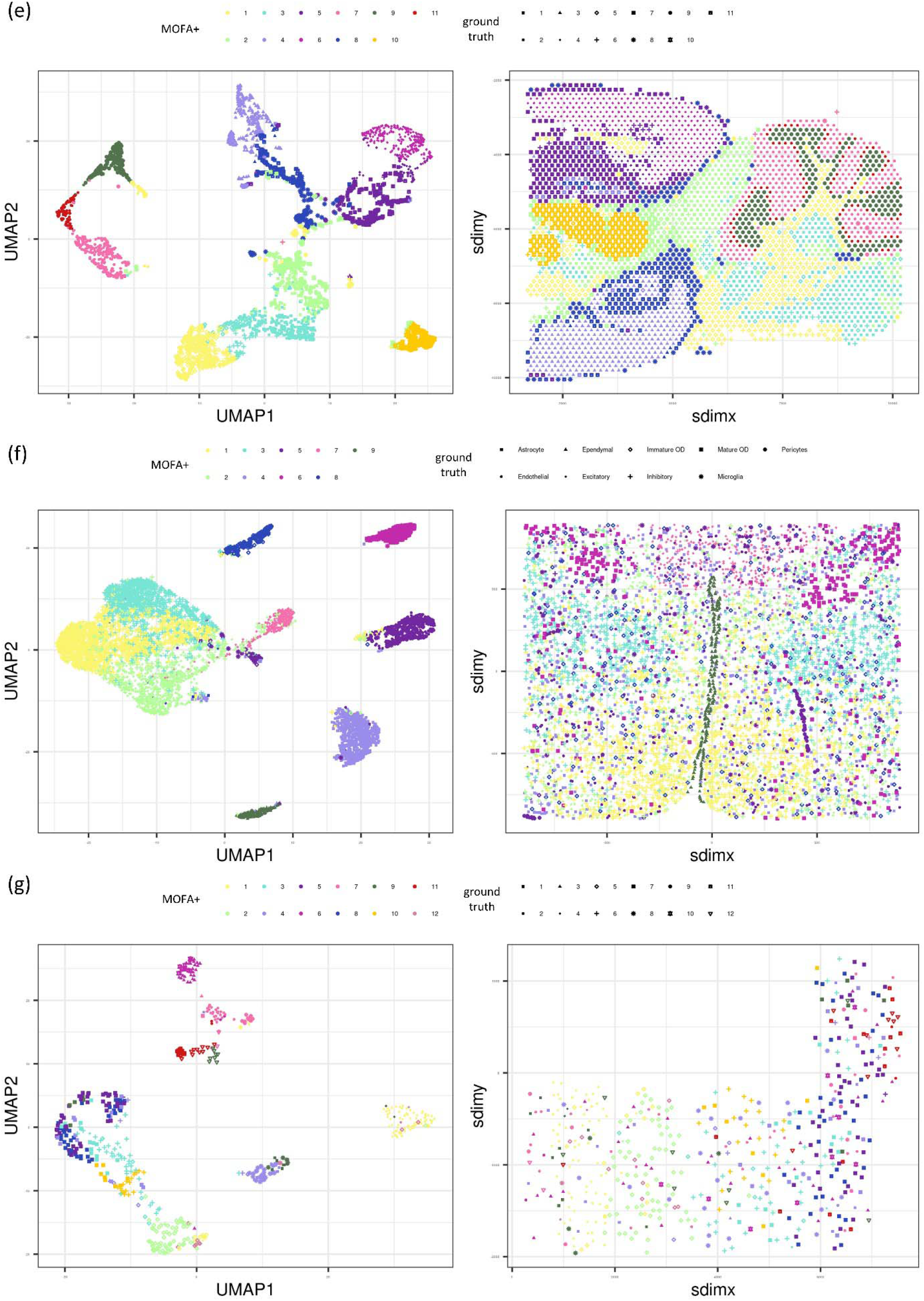
Performance comparisons of different data integration methods on real spatial transcriptomics datasets, in three representative platforms including merFISH, SeqFISH+, and Visium, across three different spatially variable genes selection methods including Giotto, spatialDE and SPARK. (a-b) Results on AMI, shown as (a) AMI boxplot of over all 19 real datasets, including two-sided paired Wilcoxon test results on the null hypothesis that the performance of the integration method is the same as the performance of solely SV genes. *: p-value < 0.05, ** : p-value < 0.01, ***: p-value < 0.001; (b) AMI ranking of different integration methods for all real datasets; (c-d) Results on VN, shown as (c) VN boxplot over all 19 real datasets, including two-sided paired Wilcox test results on the null hypothesis that the performance of the integration method is the same as the performance of solely SV genes. *: p-value < 0.05; (d) VN ranking of different integration methods for all real datasets; (e-g) comparison of MOFA+ results and ground truth labels visualized in the UMAP and spatial context for (e) the Visium Mouse Brain Sagittal Posterior dataset, (f) the merFISH Mouse Hypothalamus dataset (layer −240), and (g) the seqFISH+ Mouse Somatosensory Cortex dataset.

Since spatial transcriptomics data measure gene expression *in situ*, we also evaluated the clustering performance of each integration method with respect to the spatial distribution of the clustering labels. Towards this, we derived spatial variation norm (VN), a unique metric to fulfil this purpose (see **Methods**). The overall trends of these different integration methods are similar to those measured by AMI, although less pronounced (**Fig. 2c-d, Supplementary Fig. 2a-c**). There is no single best method for all datasets, however, concatenation has the highest average VN value and is significantly better (p-value = 0.036) than clustering solely based on SV genes detected by SPARK method (**Fig. 2c**). MOFA+ show comparable VN values to the baseline condition. Seurat v4 and scVI yield slightly worse median VN value and wider variations. Again, on average, CIMLR and SNF underperform the baseline clustering performance of using SV genes (**Fig. 2c**). We observe again technology-dependent performance patterns (**Fig. 2c-d, Supplementary Fig. 2a-c**). Seurat v4 appears to yield better rankings of VN in non-single-cell resolution Visium datasets, but worse rankings in datasets of single cell resolutions. The overall consistent trends between VN and AMI further confirms that both spatial and quantitative features are useful in cell type clustering, when the proper integration methods are used.

Additionally, we further studied the effectiveness of integration methods by visualizing the clustering and ground truth labels over spatial coordinates and two-dimensional UMAP representation based on principal components of all genes (**Fig. 2e-f**). The example results of MOFA+ for three datasets (Visium Mouse Brain Sagittal Posterior, merFISH Mouse Hypothalamus (layer −240) and seqFISH+ Mouse Somatosensory Cortex data) show that the integrated features can delineate clusters clearly, including edge cases where different clusters are not well separated on the two-dimensional UMAP representation. The clustering results exhibit clear, non-random spatial structures, further demonstrating the merit of integrating HV and SV genes in with respect to improved clustering performance.

### Clustering Accuracy on Simulated Spatial Transcriptomics Datasets

It is possible that the ground truth labels of real datasets are subject to biases from the genes and references that were used to obtain the annotations. Therefore, we also performed a simulation study where we could generate the dataset along with unambiguous ground truth labels. To simulate realistic transcriptomics data, we used the gene expression, spatial coordinates, and ground truth cell types of the seqFISH+ Mouse Somatosensory Cortex dataset as our reference [16]. We chose this dataset as the base for simulations, given its mild condition: none of the data integration methods yielded outlier values of the metrics (AMI and VN) in this dataset, among the 19 datasets. We aimed to explore how the level of spatial distinction impacted the integration and clustering performance of the methods. A gene’s spatial expression is defined as its expression pattern in the context of the spatial distribution of the cells (or spots); the more distinct the spatial pattern, the more spatially expressed the gene is. We used the spatial pattern of cell types in the ground truth to closely approximate the actual spatial transcriptomics data. We varied the level of the spatial distinction profile of SV genes by simulating new spatial patterns for these genes (**Fig. 3**). The level of spatial distinction for the new spatial pattern is controlled by a probability parameter *p* (see **Methods**). We varied the values of probability *p* from 0.6 to 0.9 and created five random repetitions for each value. We then evaluated how the probability parameter *p* affects the methods’ performance using the same pipeline (**Fig. 1**) as the previous real datasets.

**Figure 3.**
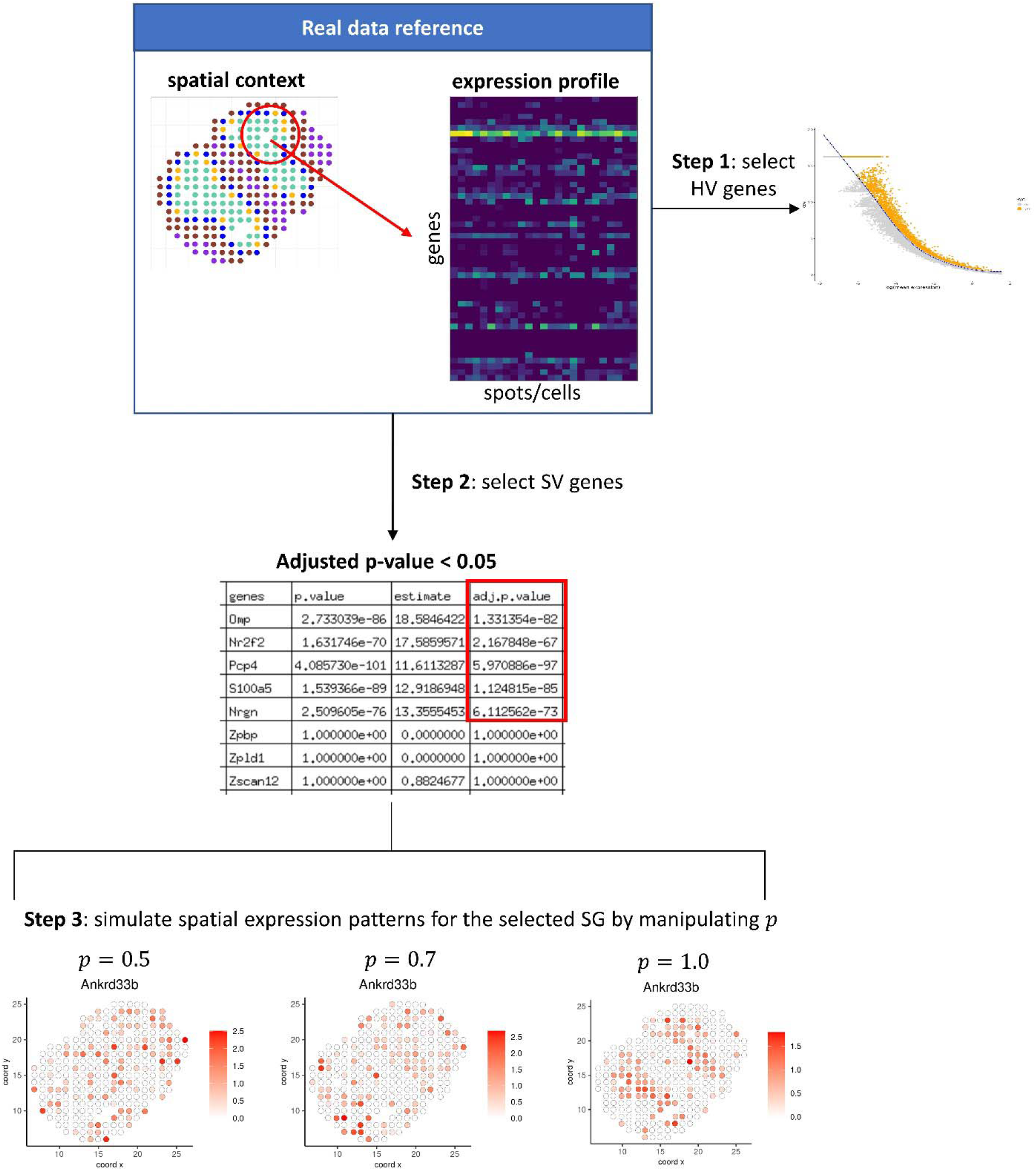
The design of simulation workflow based on the real dataset. The workflow consists of three major steps. Step 1: select HV genes. Step 2: select SV genes whose adjusted p-values are below 0.05. Step 3: simulate new spatial expression patterns for the selected SV genes.

As with real datasets analysis, we evaluated the performances of clustering results from different integration methods, compared to those based on the SV genes (**Fig. 4**). With respect to AMI, all methods except SNF and Seurat v4, consistently yield significantly higher values compared to the baseline condition of using SV genes (**Fig. 4a**). Moreover, as the level of spatial distinction increases, the AMI values of all methods continue to improve, confirming that the level of spatial distinction certainty is informative of cell type clustering (**Fig.4a**). Under varying levels of spatial distinction, MOFA+ has the best and the most stable spatial omics clustering performance (AMI) among all the integration methods. Other methods such as concatenation, CIMLR and scVI also do well. Note that the advantage of MOFA+ over the rest of the methods is more apparent when the spatial distinction is relatively small. This suggests that MOFA+ is able to effectively integrate HV and SV genes even with low certainty. Again SNF has relatively worse stability over different levels of spatial distinction.

**Figure 4.**
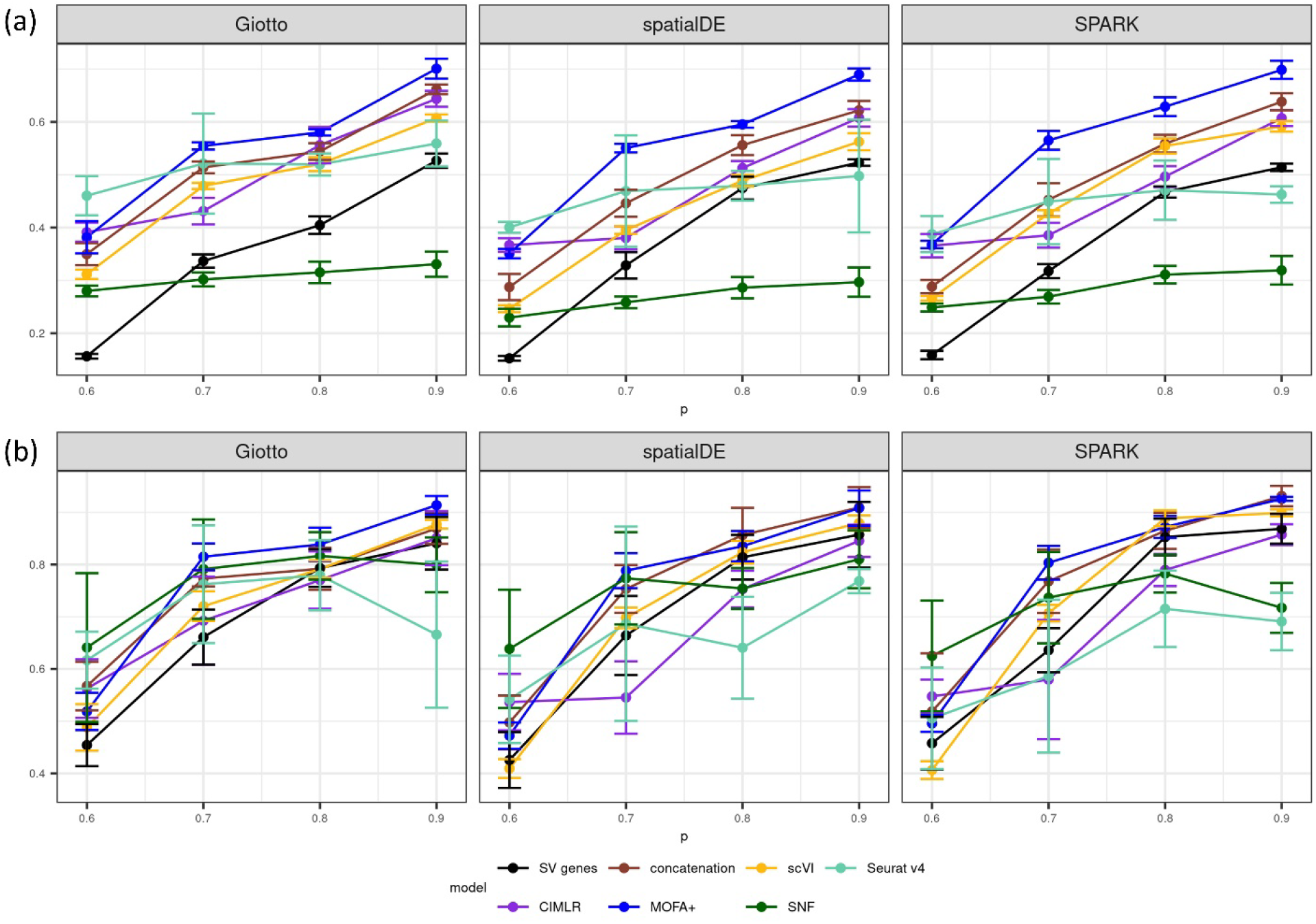
Clustering performance comparisons of different data integration methods on simulated spatial transcriptomics datasets, by varying probability (*p*) of spatial clustering certainty (0.6, 0.7, 0.8, 0.9), for different spatially variable gene selection methods including Giotto, spatialDE and SPARK. Shown are metrics using (a) AMI and (b) VN. Five repetitions are run in each condition, and the average scores are displayed, with standard deviations.

For spatial clustering evaluation using VN, we observe an overall increasing accuracy trend as the level of spatial distinction increases for most integration methods (**Fig.4b**). MOFA+ and concatenation have the best and most stable spatial clustering performance among all the integration methods. Both methods consistently outperform spatial clustering solely based on SV genes, over varying degrees of spatial distinction. scVI also consistently outperforms spatial clustering based on SV genes, however the advantage is not as apparent. CIMLR, SNF and Seurat v4 have lower VN values compared to those using solely SV genes, and Seurat v4 has the worst stability over different levels of spatial distinction.

### Effect on Downstream Differential Expression

Clustering analysis is usually an important step for other downstream analyses. In order to further assess the downstream effect of clustering through integrating SV and HV genes, we performed differential expression analysis. For this we used the differential expression analysis results based on the ground truth labels as the references. We tested how many original markers can be recovered based on the integration cluster labels. We evaluated the downstream differential expression analysis performance using the F-1 score (see **Methods**), in comparison with that of the clustering based solely on SV genes.

For the 19 real datasets, scVI had the highest median F-1 score and significantly better than the baseline performance based solely on SV genes (p-value = 0.036, 0.520, and 0.418 for Giotto, spatialDE, and SPARK). scVI’s performance is followed closely by MOFA+ and concatenation. Seurat v4 achieved worse F-1 scores comparable to that of SV genes, but the disadvantage is generally not significant. SNF and CIMLR yielded significantly worse F-1 scores than the baseline, both with p-values less than 0.01 (**Fig. 5a**). The specific values and rankings of the F-1 scores again present technology dependent patterns, similar to those observed by AMI and VN. Seurat v4 obtains better rankings of F-1 scores in non-single-cell-resolution datasets than in single-cell resolution datasets (**Fig. 5a-b, Supplementary Fig. 3a-c**).

**Figure 5.**
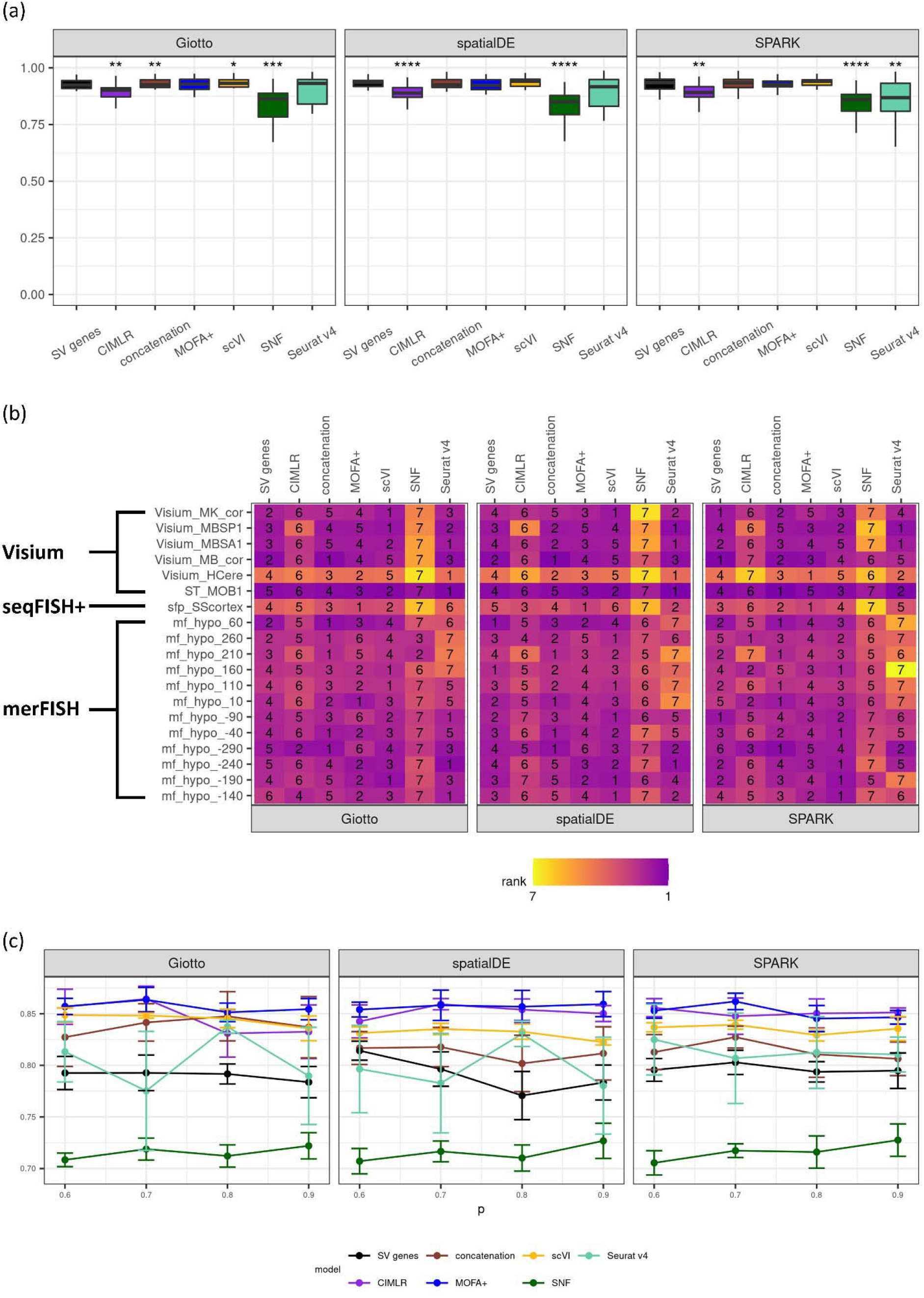
Effect of integration methods on downstream differential expression results measured by F-1 scores, for different spatially variable gene selection methods including Giotto, spatialDE and SPARK. (a-b) Results on real datasets, in three representative platforms, including merFISH, SeqFISH+, and Visium, shown as (a) F-1 score boxplot over all 19 real datasets, including twosided paired Wilcox test results on the null hypothesis that the performance of the integration method is the same as the performance of solely SV genes. *: p-value < 0.05, ** : p-value < 0.01, ***: p-value < 0.001, ****: p-value < 0.0001; (b) F-1 score ranking for all real datasets; (c) Results on the same simulation datasets as in Figure 4.

We performed similar downstream differential expression analysis for the simulated datasets (**Fig. 5c**). MOFA+ has the best and most consistent performance across different SV genes detection methods and varying levels of spatial distinction. Besides MOFA+, CIMLR, scVI and concatenation also outperform the baseline of using SV genes alone. SNF shows significantly worse F-1 scores than the baseline condition. The F-1 score of Seurat v4 is relatively unstable compared to other integration methods. It is generally worse than SV genes at lower levels of spatial distinction, but improves when the level of spatial distinction is stronger.

### Runtime and Memory Usage

Runtime efficiency and memory usage are two important practical measures for computational methods. Therefore, we used real Human Cerebellum dataset by Visium platform for testing. We compared the runtime and memory usage of the six integration methods by randomly subsetting the number of cells from 500, 1000, 1500, 3000, to 5000 and the number of features from 500, 1000, 3000, 5000, to 10000. All the integration methods were run on the Garmire Lab server, which operated on a Linux operating system (Intel(R) Xeon(R) Gold 6148 CPU @ 2.40GHz, 219 GB RAM, 16 GB Nvidia GPU). MOFA+, scVI and SNF were run on GPU nodes and concatenation, CIMLR and Seurat v4 ran on regular nodes.

Increasing the number of cells has a more significant impact on both runtime efficiency and memory usage than increasing the number of features (**Fig. 6**). For running time measurements, straightforward concatenation is the most time-efficient method among all, as expected (**Fig. 6 a, c**). The other iterative computational methods such as Seurat v4, CIMLR, SNF, MOFA+ and scVI have an ascending order for computing time. Increasingly longer runtimes are needed, as the number of cells or the number of features increases (**Fig.6 a, c**). For memory usage, concatenation is the most memory-efficient method over varying numbers of cells and features. On the other hand, CIMLR uses the most memory, which increases significantly more than the rest of the methods as the number of cells increases. This could be attributed to CIMLR’s adaptation of parallel computation. scVI uses the most memory as the number of features increases; it is closely followed by MOFA+ and CIMLR.

**Figure 6.**
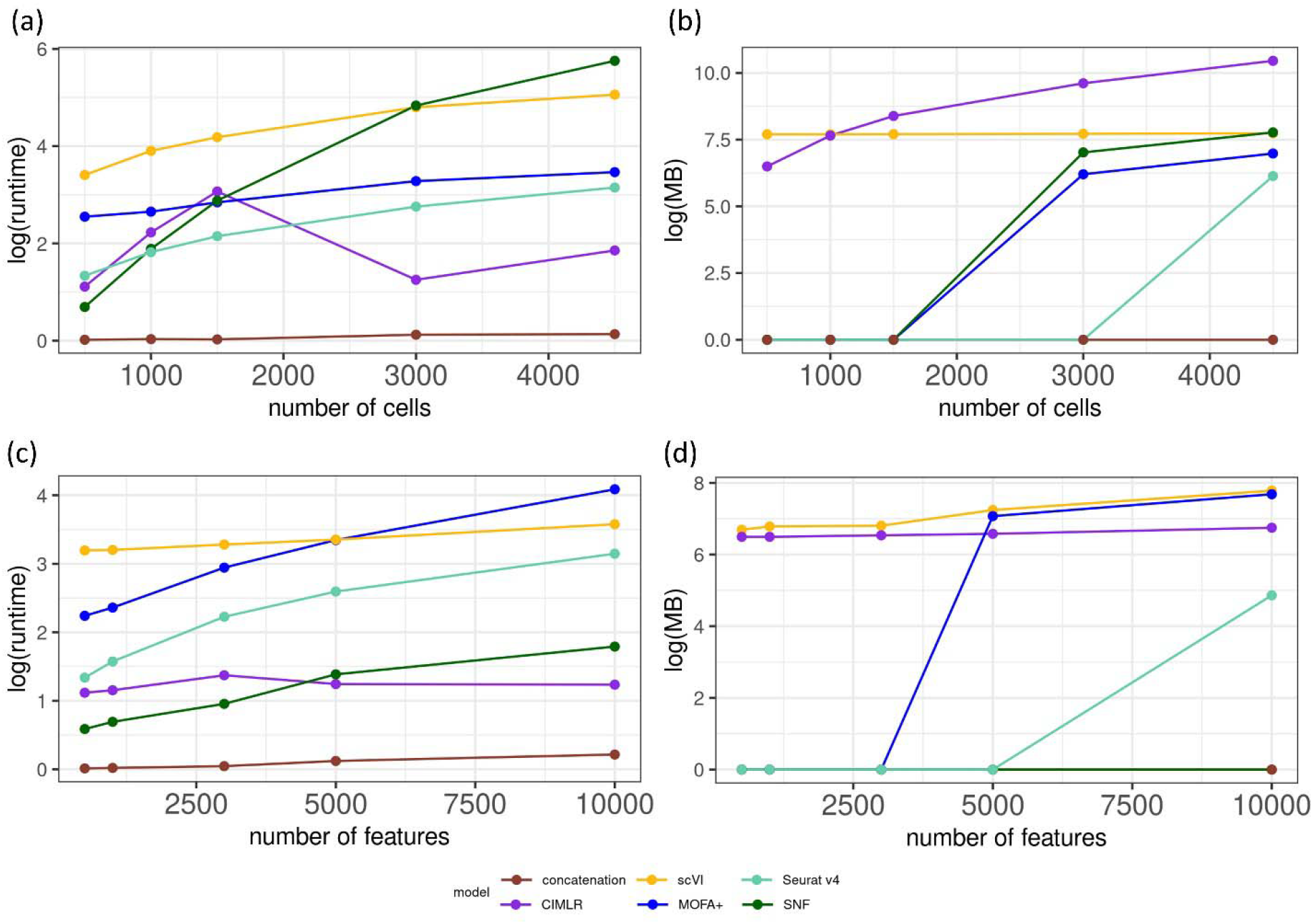
Comparison of running time and memory usage of integration methods using the Visium Human Cerebellum dataset. (a-b) The practical performance relative to the number of cells, measured by (a) running time, and (b) memory usage. (c-d) The practical performance relative to the number of features, measured by (c) running time, and (d) memory usage.

## Discussion

The advent of spatial transcriptomics technologies gives us the potential to create a more comprehensive map of biological systems. Relative to single cell RNA-Seq technologies, the addition of spatial information has the potential to help discover novel SV markers. Previously, SV markers have been used for identifying clusters in spatial transcriptomics data; SV genes that share similar spatial expression patterns can also be used to define cell types as well as relate cell type composition to tissue structure [18–21]. However, these SV markers represent genes whose expression pattern across space is non-random, which is a different category from those HV gene markers detected by quantitative variabilities conventionally [37–40]. Questions remained: (1) if SV gene based clustering can be improved by integrating additional HV genes, which are normally used in single cell RNA-Seq and bulk RNA-Seq analysis for clustering; (2) if integration of SV and HV genes can improve clustering results in spatial transcriptomics data, which computational method(s) are suitable. Since there are no current gold-standard pipelines for clustering spatial transcriptomics data specifically, our study fills the niche by conducting a systematic benchmarking study. We have confirmed that combining these two types of markers can improve the clustering results, with appropriate integration methods.

**Fig. 7** shows an overview of the spatial omics clustering accuracy (by AMI), spatial clustering accuracy (by VN), downstream differential analysis performance (by F-1 score), and runtime and memory usage for 19 real datasets across different technology platforms and 20 simulated datasets across different spatial distinction levels. We recommend MOFA+ for general spatial transcriptomics datasets; straightforward concatenation is also a good approach especially considering the computational cost (running time and memory use). However, simulation results show concatenation is more sensitive to the level of spatial distinction than MOFA+. Technology-specific patterns show that Seurat v4 performs better for integrating HV and SV genes of non-single-cell resolution datasets obtained from the Visium platform. SNF generally yields worse results than the baseline condition using solely on SV genes. Therefore, the current implementation of SNF is not suitable for integrating SV and HV genes in spatial transcriptomics data.

**Figure 7.**
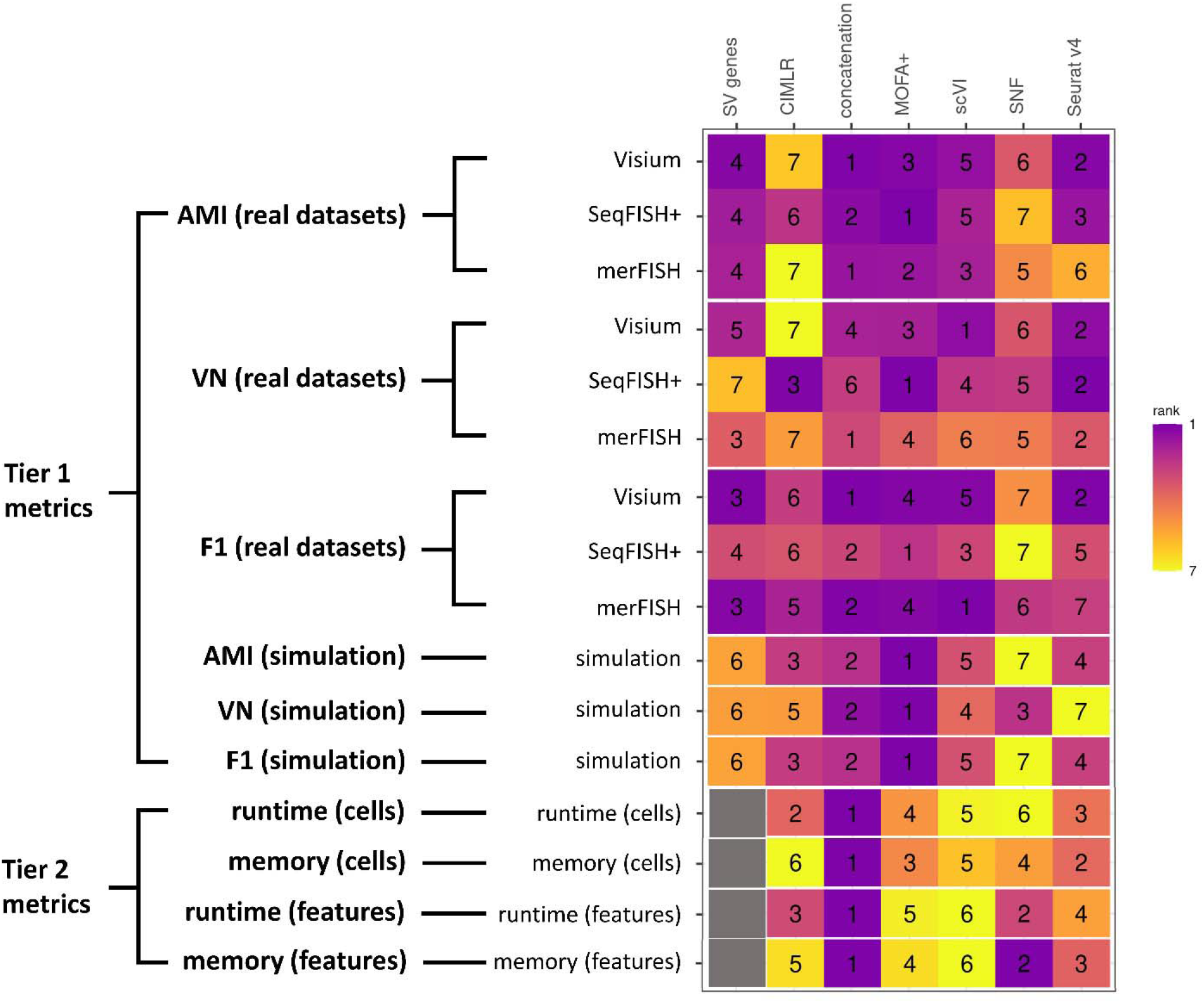
Heatmap summary of metrics ranking of integration methods over real and simulated datasets, by Tier 1 metrics: AMI, VN, F-1 statistics of downstream differential expression analysis, and Tier 2 metrics: runtime and memory usage.

The quality of clustering evaluation above is dependent on the ground truth labels. The ground truth labels of 18 out of 19 datasets are derived using all the genes, except for the one seqFISH+ dataset whose ground truth labels are based on highly variable genes (see **Table 2**). Therefore, the ground truth labels of the real datasets are generally unbiased toward highly variable genes. Moreover, the 20 simulated datasets also support the general conclusion from the real datasets, in that MOFA+ and concatenation approaches yield better results over other integration methods in general.

MOFA+ belongs to a class of matrix-factorization-based methods. Our evaluation shows that it may improve clustering results in integrating HV and SV genes, as well as subsequent downstream differential expression analyses. This class of models was previously shown quite effective in utilizing information from multiple sets of genomics features and extracting useful information for integration [29,41,42]. In particular, MOFA+ decomposes each component dataset as a product of a shared dimension-reduced latent space and a dataset-specific weight matrix. MOFA+ model can learn the shared dimensionally-reduced latent space by borrowing information from each component dataset, thus allowing for effective integration (see **Method**). Despite the additional runtime and memory usage, MOFA+ also provides an additional advantage over simple concatenation, as the shared latent space learned by MOFA+ can provide more insight into spatial transcriptomics features. Dimension reduction also appears to be an important factor in successful integration of features. Both Seurat v4 and SNF are based on building joint networks, but Seurat v4 has significantly better performance than SNF. Seurat v4 performs dimension reduction before building the networks whereas SNF attempts to directly integrate networks built on data not dimensionally reduced. Though slightly better than SNF, CIMLR generally performs worse than the other four methods. This could be attributed to the Gaussian kernel assumption that CIMLR makes (see **Method**). Such an assumption might oversimplify the complexities of spatial transcriptomics data.

## Conclusions

In general, we recommend MOFA+ for integration of quantitatively variable and spatially variable genes in spatial transcriptomics data. We also stress the need for methods to be developed in this area which are further complicated by different spatial transcriptomics technology platforms.

## Methods

### Data Preprocessing

For all the computational methods including extracting HV and SV genes, we preprocessed the data by first filtering the raw gene expression dataset. The expression threshold was set to 1. The minimum number of cells a gene needs to be expressed in was set to 3 and the minimum number of expressed genes in a cell was set to at least 5% of the total number of genes. Such values were chosen to filter out unexpressed genes or cells without over-processing the data. The SV genes method SPARK requires a higher threshold for filtering genes in order to avoid internal computational errors. Therefore, we raised the gene filtering threshold to 0.75% of the total number of cells for SPARK analyses. As was shown in the benchmarking results, this discrepancy did not cause additional inconsistencies.

Computational methods such as straightforward concatenation, MOFA+, SNF and CIMLR require normalized gene expression data. For these methods, the filtered raw gene expression data was normalized via log-normalization, and the cells and genes were scaled by Giotto’s [21] default scale factor of 6000. We skipped normalization for Seurat v4 and scVI, as they require raw data.

### Benchmark Analysis Parameters

We extracted the HV genes and the SV genes based on normalized expression data. Among the computational methods, MOFA+, scVI, Seurat v4 and CIMLR have inherent dimension reduction steps (see Supplementary Table 1 for details) which extract combined features on a lower dimension. For the rest of the methods that do not have inherent dimension reduction steps, we performed dimension reduction using PCA (Principal Component Analysis). The dimension of the extracted features was set to be the same across all methods for a fair comparison. The optimal number of features were determined empirically by comparing the general clustering performance across different values of reduced dimensions. We used Leiden clustering for all methods [43] except for CIMLR, which had its own built-in clustering step. For a fair comparison of evaluation metrics, we repeatedly performed Leiden clustering, tuning the resolution and number of nearest neighbors parameters for each computational method until we got the same number of clusters as there were ground truth clusters.

### Selection of HV and SV genes

We used HV (highly variable) genes as features whose expression was informative of how the data clustered. We extracted HV genes using the R package *Giotto’s calculateHVG* function [21], The function fits a LOESS regression model with each gene’s log mean expression as the independent variable and the coefficient of variance as the dependent variable. Genes with a coefficient of variance that was higher than their predicted value past an arbitrary threshold were considered to be highly variable. We set the threshold value to the default: 0.1.

We used SV genes as features whose expression was informative of how clusters were distributed spatially. We explored three popular SV genes methods: Giotto, spatialDE and SPARK. For Giotto, we extracted the SV genes using the R package *Giotto’s binSpect* function [21]. The function binarized each gene’s expression value using the K-means method (number of centers set to 2). The function then constructed a spatial connectivity network based on the cells/spots’ spatial coordinates using Delaunay’s triangulation [44]. A contingency table was generated based on the connectivity of the spatial network and the binarized gene expression. The function performed the Fisher’s test and generated a spatial expression score for each gene with higher values indicating a more distinct spatial expression pattern.

For fair comparison across SV genes methods, we used the number of SV genes discovered by Giotto as the imposed number of SV genes by spatialDE and SPARK. For both spatialDE and SPARK, we used their wrapper function in the R package Giotto. spatialDE models the spatial gene expression data using a spatial linear regression model based on Gaussian Processes. The spatial significance of a specific gene is determined through a likelihood-ratio-type hypothesis test on the spatial effects term in the model. Similarly, SPARK can be considered as an extended, generalized version of spatialDE model that can directly analyze raw spatial transcriptomics data.

Since HV genes and SV genes usually overlap, to avoid double counting the overlapped genes in integration, we defined HV genes as the genes that are uniquely highly variable in the integration step. SV genes are those that are uniquely spatially variable and those that overlap. Suppose there are *g*_1_ unique HV genes and the optimal number of reduced dimensions is *k*. We denote the subset of normalized HV genes as *E*_1_ (*g*_1_ genes × *m* cells). We performed PCA on *E*_1_ and selected the top *k* principal components, represented by a *k* × *m* matrix denoted as *W_HV_*. We used *W_HV_* for further clustering analysis. Similarly, suppose there are *g*_2_ unique SV genes. We denoted the normalized SV genes as *E*_2_ (*g*_2_ genes × *m* cells). We performed PCA on *E*_2_ and selected the top *k* principal components, represented by a *k* × *m* matrix denoted as *W_SV_*. We used *W_SV_* for further clustering analysis.

### Methods in comparison

#### Simple concatenation model

we concatenated the normalized HV genes *E*_1_ and the normalized SV genes *E*_2_, represented by a (*g*_1_ + *g*_2_) × *m* matrix denoted as *E*. We performed PCA on the concatenated dataset *E* and took the top *k* principal components for further clustering analysis.

#### CIMLR (Cancer Integration via Multikernel Learning) model [27]

we adapted CIMLR, which was originally developed to integrate multi-omics cancer data, to integrate normalized HV genes *E*_1_ and normalized SV genes *E*_2_. CILMR constructed a group of Gaussian kernels based on *E*_1_ and *E*_2_ separately. CIMLR then computed the similarity matrix between the two datasets by combining the gaussian kernels. Given the number of clusters, dimension reduction and clustering via k-means clustering were then performed on the similarity matrix.

#### MOFA+ (Multi-Omics Factor Analysis v2) model [25]

MOFA+ is a statistical framework for integrating multi-omics single-cell data based on representing the original dataset in lower dimensions through matrix factorization infused with a Bayesian technique for enhancing model sparsity. We adapted MOFA+ to integrate HV genes and SV genes. We used the normalized HV genes *E*_1_ and the normalized SV genes *E*_2_ as input. MOFA+ aims to reconstruct *E_i_* using lower dimensional matrices *W_MOFA+_* (*m* × *k*) and *U_i_* (*g_i_* × *k*): (*ϵ_i_* represents random noise) (Eq.2). *W_MOFA+_* represents the low-dimensional factors common across all datasets and *U_i_* represents each feature’s association score with each factor. We used *W_MOFA+_* for further clustering analysis.

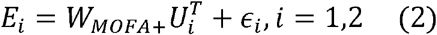

#### scVI (Single-Cell Variational Inference) model [23]

scVI constructs latent space representations of cells using a variational autoencoder. The *encoder* neural network maps gene expression values onto parameters of the probability distribution from which latent variables are sampled, while the *decoder* neural network takes as input such latent variables and aims to reproduce the original input, by producing the parameters of the ZINB (zero-inflated negative binomial) distribution modelling the gene expression values. The model also uses neural networks to model dropout patterns and adds scales to account for differences in gene expression between different cells. We concatenated the raw HV genes and the raw SV genes and fed them as input to the scVI model. We used scVI with a single hidden layer in an encoder and optimized the dimension of the hidden layer, and the dropout rate through a grid search, optimizing for the minimum reconstruction error.

#### SNF (Similarity Network Fusion) model [28]

we adapted the SNF algorithm to construct graph representations of normalized HV genes *E*_1_ and normalized SV genes *E*_2_ and merged them to construct a coherent picture. Each node on such graphs corresponds to different cells and the edge weights correspond to the degree of similarity (correlation) between them when separately considering highly variable and SV genes. The graphs were fused through iterative updates, inspired by diffusion processes, which gradually bring the graphs closer together. We performed clustering analysis on the integrated similarity graph.

#### Seurat v4 (Weighed Nearest Network) model [24]

we adapted Seurat v4 to compute cell-specific weights separately for the raw HV genes and the raw SV genes. Such weights reflected the “usefulness” of information from HV genes and SV genes. Seurat v4 integrated the HV genes and SV genes while leveraging their constructed weights so the integrated data reflected the information of its component datasets. Nearest neighbor graph construction and clustering were then performed on the integrated data.

### Ground truths for real data sets

The quality of the ground truth labels are essential to the evaluation of methods’ performance. We had complete control over the ground truth labels of the simulated datasets as they were arbitrarily generated. For the real datasets, we obtained ground truth labels from the original studies whenever possible. For the Mouse Olfactory Bulb dataset [16] generated by the technology that was later named Visium, we were able to recreate the cell type assignments based on the original supplementary materials. For the rest of the five Visium datasets, we used the clustering labels provided by 10X [32–36]. For the Mouse Somatosensory Cortex dataset by SeqFISH+ [14], we used the clustering labels generated by *Giotto* package [21]. For the merFISH datasets, we used the cell type annotations in the original report [10].

### Simulation Study

Using the seqFISH+ Mouse Somatosensory Cortex dataset as reference, we used the original unique HV genes of the real dataset as the HV genes in this simulation study. We selected SV genes whose adjusted p-values are below 0.05. To ensure the simulated spatial expression patterns are meaningful with respect to identifying the ground truth cell types, we matched different SV genes to the ground truth cell type whose spatial expression pattern had the highest level of concordance.

We adapted *Giotto’s* [21] *simulateOneGenePatternGiottoObject* function to simulate spatial expression profiles for each individual selected spatially variable gene. We used the spatial patterns of cell types in the ground truth to guide customized simulations for spatial gene expression patterns. Given a customized pattern that contains *n* cells, with probability *p*, the spots where the gene is highly expressed will fall into the customized pattern in the simulated dataset. When *p* = 0.5, the cells will fall into the customized pattern completely randomly; when *p* = 1.0, the cells will fall into the customized pattern with absolute certainty. We tested four different values for spatial probability *p* (0.6, 0.7, 0.8, 0.9) to represent different noise levels in clustering labels, and repeated the experiment 5 times for each value of *p*.

### Hypothesis Tests

We performed non-parametric paired Wilcox tests to further investigate the efficacy of the performance of the integration methods in comparison to the baseline performance of solely SV genes. For each metric (AMI for omics clustering, VN for spatial clustering, F-1 for downstream differential expression analysis), we performed the paired Wilcox test on each pair of SV genes metrics and integration method metrics. The null hypothesis is that the metric performance of solely SV genes comes from the same distribution as the performance of the integration method. The methods whose corresponding p-values are less than 0.05 were considered to perform statistically significantly different from the baseline SV genes.

### Evaluation Metrics

#### Adjusted Mutual Information (AMI)

To evaluate general accuracy of clustering in spatial transcriptomics data, we computed the Adjusted Mutual Information (AMI) [45] of each set of clustering results compared with the ground truth. AMI measures the level of concordance between two sets of labels and is widely used for measuring clustering accuracy in scRNA-Seq datasets with no spatial context [46–48]. AMI is bounded between 0 and 1, with higher values indicating better clustering performance.

#### Variation Norm (VN)

To evaluate the clustering results accuracy in the spatial context, we used a metric based on the total variation norm denoted as VN [49–52]. The larger VN is, the more accurate the spatial clustering is compared to the ground truth. We denote the set of clustering labels of a certain method as *u*\* and the set of ground truth labels as *u*. For each set of labels, we computed the average pairwise Euclidean distance between the cells or spots within each cluster. We denote the density functions of such group-wise average distance as *q*(*s*|*u*\*) for the clustering labels *u*\* and *q*(*s*|*u*) for the clustering labels *u*, respectively. Thus, comparison of *q*(*s*|*u*\*) and *q*(*s*|*u*) allows for evaluation of clustering accuracy while taking into account the relative spatial structure of the cells / spots. We computed the total absolute error *IAE*(*u*\*) as

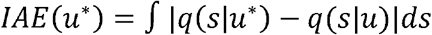

The accuracy measure VN is defined as

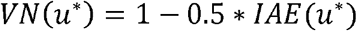

#### F-1-score in differential expression

we performed traditional quantitative downstream differential expression analysis over the clustering results in a one-vs-all manner using the marker gene detection method “*Gini*” in Giotto package [21]. We used the marker gene detection results based on the ground truth clustering labels as reference. We compared the marker genes detected based on the integration method vs. those in the reference. Considering the marker gene detection as a binary classification problem, we counted the number of true positives (TP), false positives (FP) and false negatives (FN). We then computed the F-1 score. The larger the F-1 score is, the more accurate the prediction.

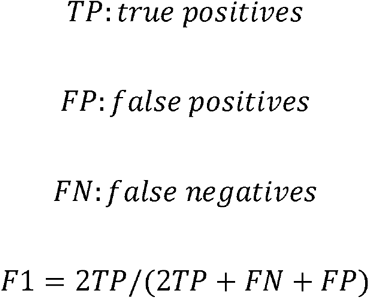

## Declarations

### Ethics approval and consent to participate

All data utilized in this study has been previously published and is publicly available. The research performed in this study conformed to the principles of the Helsinki Declaration

### Consent for publication

Not Applicable

### Availability of data and materials

All data and code used in this benchmark study can be found in the following link https://github.com/lanagarmire/ST_benchmark

## Author Contributions

LG envisioned this project. YL developed the benchmark framework, simulation design, implemented the computational methods and evaluated the methods’ performance. SS helped implement scVI and SNF, and double checked reproducibility. BH and QH helped select the real datasets. ZJ helped implement MOFA+. JK helped develop the spatial clustering evaluation metric. All authors have read and approved the final manuscript.

## Acknowledgment

Not Applicable.

