## supplementary figures and tables for "Benchmarking Computational Integration Methods for Spatial Transcriptomics Data"

**Supplementary materials**

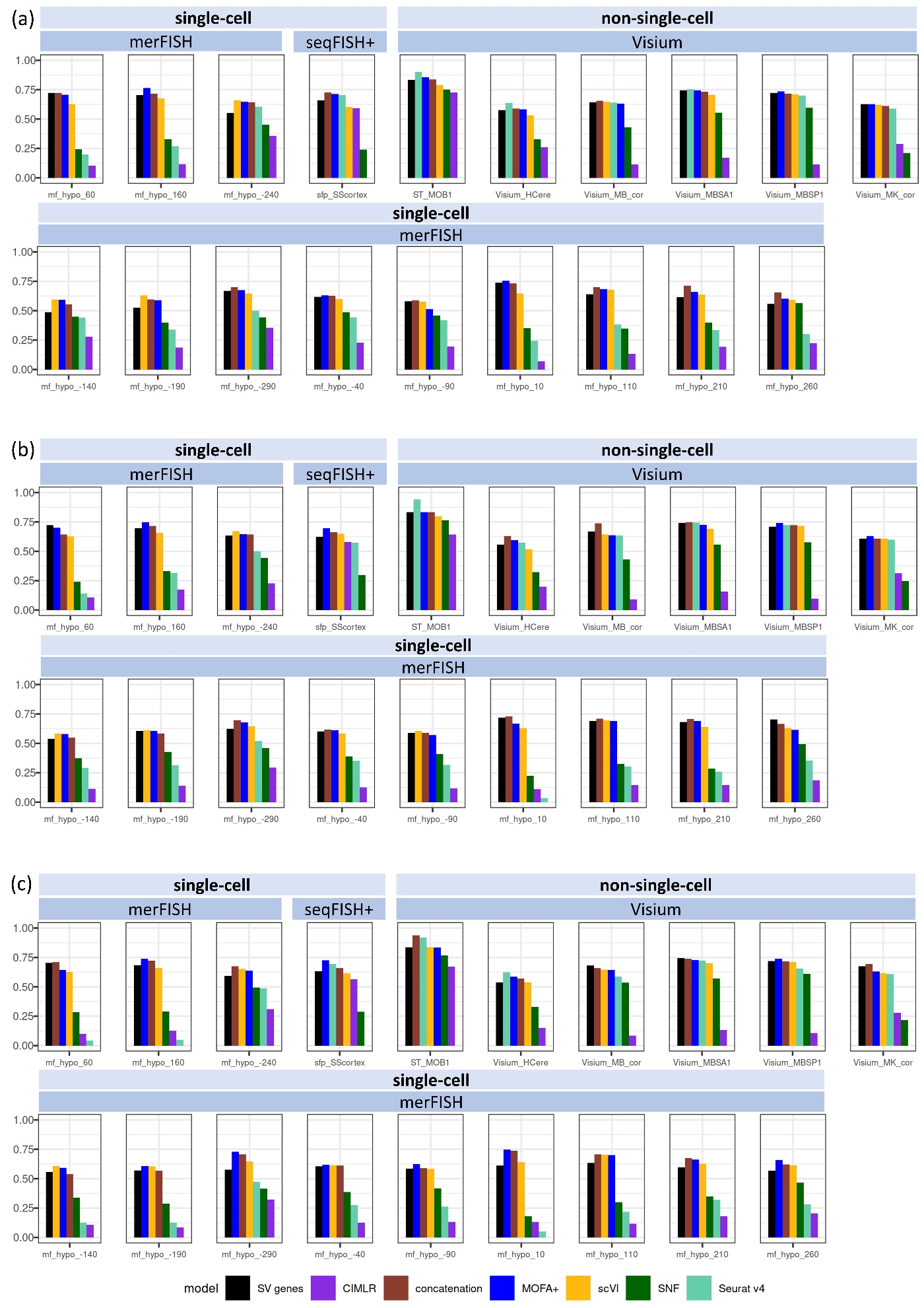

**Supplementary Figure 1.** Barplot of AMI values for the all 19 datasets for different spatially variable genes selection methods: (a) Giotto, (b) spaitalDE, and (c) SPARK.

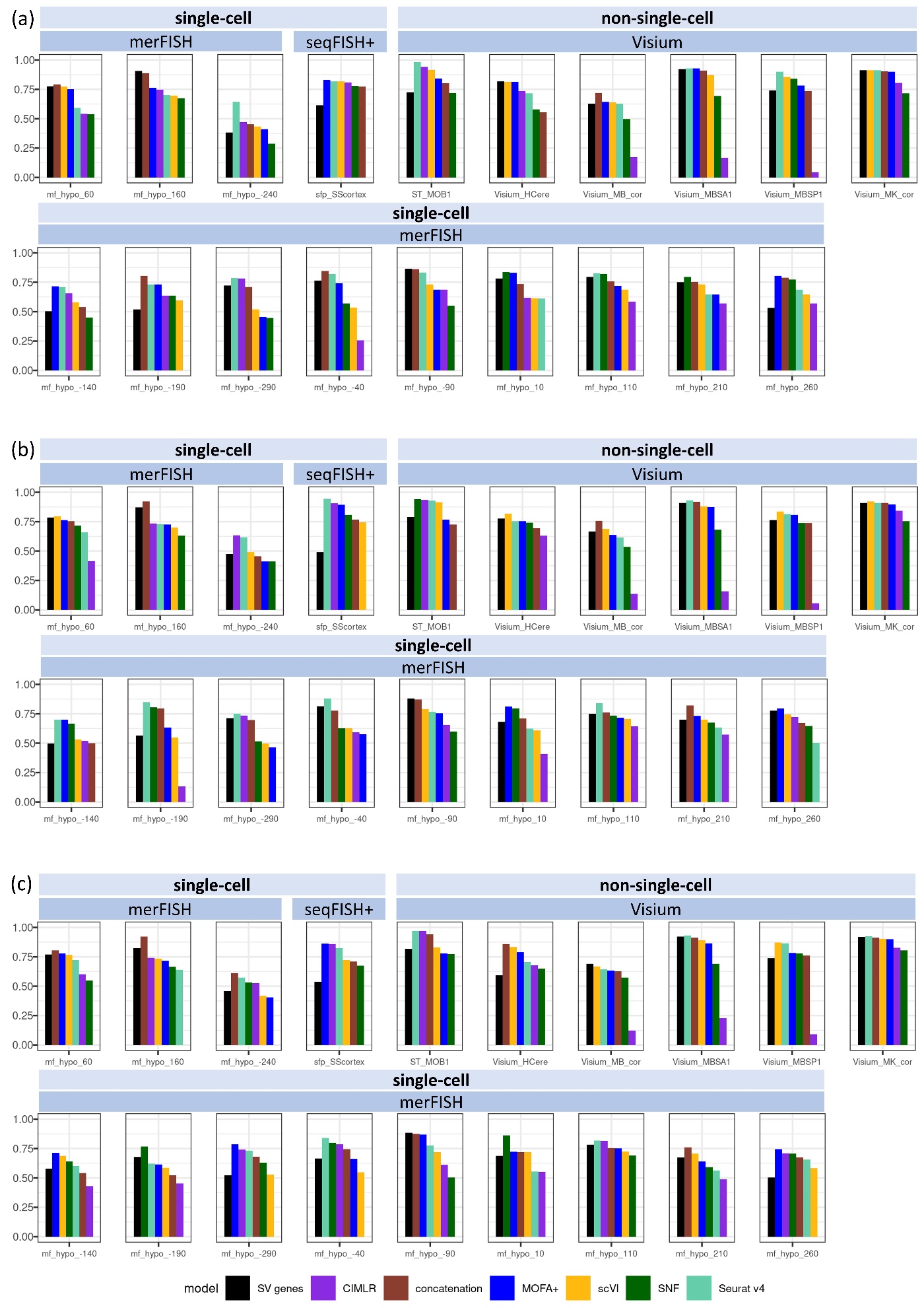

**Supplementary Figure 2.** Barplot of VN values for the all 19 datasets for different spatially variable genes selection methods: (a) Giotto, (b) spaitalDE, and (c) SPARK.

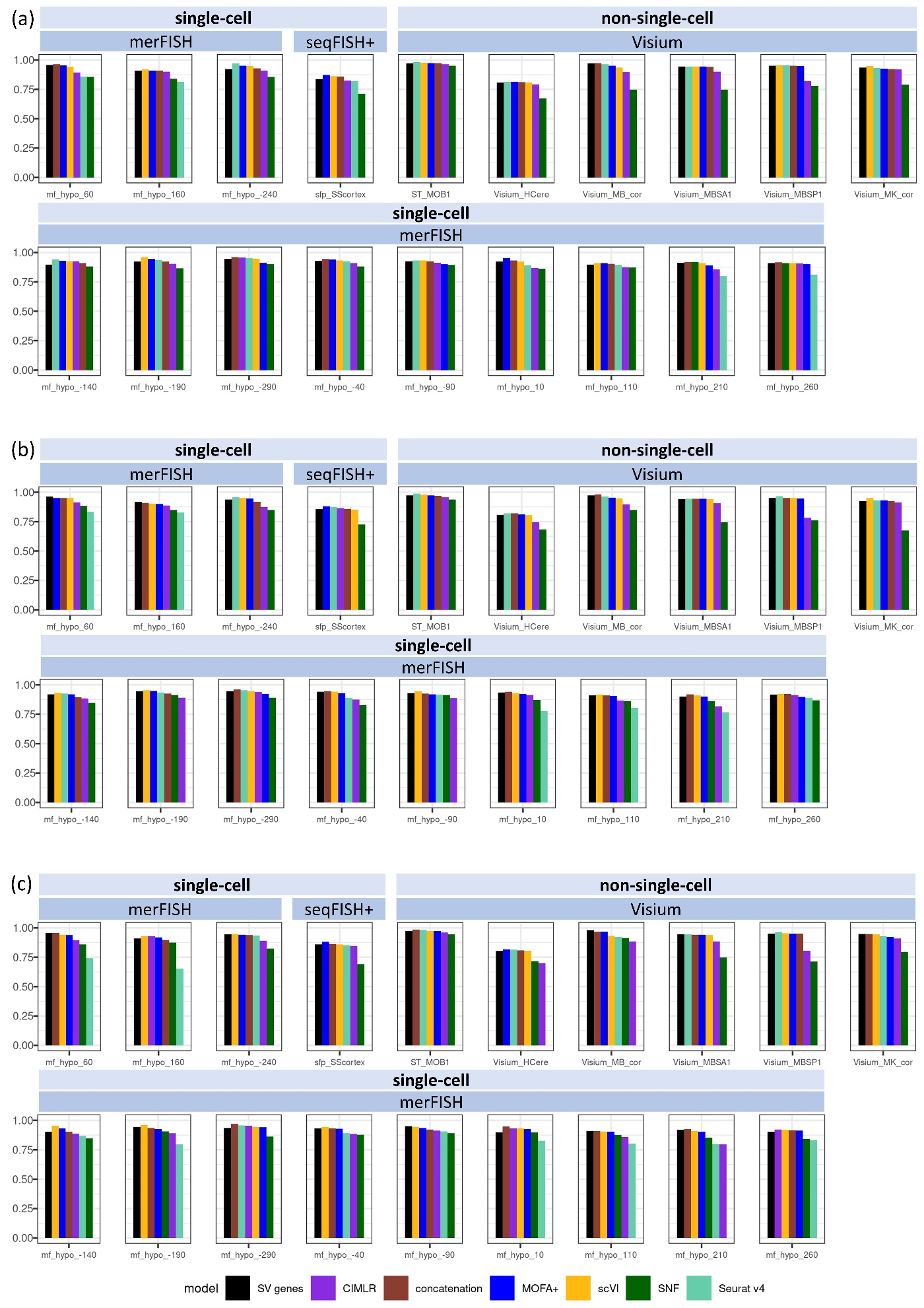

**Supplementary Figure 3.** Barplot of F-1 values for the all 19 datasets for different spatially variable genes selection methods: (a) Giotto, (b) spaitalDE, and (c) SPARK.

**Table 1**: Summary of integration methods in comparison.

| **Method** | **Working principle** | **Existence of Internal dimension reduction step** | **Required input data** |
| --- | --- | --- | --- |
| Concatenation | Straightforward concatenation | no | Normalized |
| CIMLR | Kernel-based method | yes | Normalized |
| MOFA+ | Matrix Factorization | yes | Normalized |
| scVI | Variational Autoencoder | yes | Raw |
| SNF | Similarity network based fusion method | NA | Normalized |
| Seurat v4 | Weighted Nearest Neighbor | yes | Raw |

**Table 2**: Summary of datasets and their spatial transcriptomics platforms.

| **Study** | **Genes utilized in ground truth labelling** | **Technology**  **platform** | **Resolution** | **Number of cells (or spots)** | **Number of features** | **HVG** | **SVG** | **HVG & SVG (Giotto)** | **HVG & SVG (spatialDE)** | **HVG & SVG (SPARK)** |
| --- | --- | --- | --- | --- | --- | --- | --- | --- | --- | --- |
| Mouse Olfactory Bulb [18] | All | Visium | non-single-cell | 265 | 16573 | 3938 | 1928 | 647 | 753 | 822 |
| Mouse Somatosensory Cortex [[14]](https://paperpile.com/c/xwk5cU/up8MB) | highly variable genes | SeqFISH+ | single-cell | 523 | 10000 | 1171 | 1151 | 142 | 182 | 164 |
| Human Cerebellum [[31]](https://paperpile.com/c/xwk5cU/4WDQb) | All | Visium | non-single-cell | 4992 | 23808 | 5967 | 9325 | 2256 | 3008 | 2927 |
| Mouse Kidney Coronal Section [[32]](https://paperpile.com/c/xwk5cU/cXIGV) | All | Visium | non-single-cell | 1438 | 20100 | 3888 | 7983 | 1845 | 1929 | 1855 |
| Mouse Brain Coronal Section [[33]](https://paperpile.com/c/xwk5cU/fQXCZ) | All | Visium | non-single-cell | 2702 | 21949 | 4255 | 12152 | 2503 | 2534 | 2455 |
| Mouse Brain Sagittal Anterior Section [[34]](https://paperpile.com/c/xwk5cU/mCOGA) | All | Visium | non-single-cell | 2695 | 21334 | 4184 | 12769 | 2368 | 2511 | 2503 |
| Mouse Brain Sagittal Posterior Section [[35]](https://paperpile.com/c/xwk5cU/99lBA) | All | Visium | non-single-cell | 3355 | 21363 | 4677 | 13207 | 2531 | 2505 | 2618 |
| Mouse Hypothalamus (layer -240) [[10]](https://paperpile.com/c/xwk5cU/7Ph4u) | All | merFISH | single-cell | 5543 | 157 | 81 | 88 | 47 | 49 | 50 |
| Mouse Hypothalamus (layer -190) [[10]](https://paperpile.com/c/xwk5cU/7Ph4u) | All | merFISH | single-cell | 5803 | 156 | 80 | 103 | 54 | 57 | 59 |
| Mouse Hypothalamus (layer -140) [[10]](https://paperpile.com/c/xwk5cU/7Ph4u) | All | merFISH | single-cell | 5926 | 157 | 82 | 104 | 53 | 59 | 58 |
| Mouse Hypothalamus (layer 60) [[10]](https://paperpile.com/c/xwk5cU/7Ph4u) | All | merFISH | single-cell | 5343 | 157 | 78 | 119 | 55 | 57 | 58 |
| Mouse Hypothalamus (layer -40) [[10]](https://paperpile.com/c/xwk5cU/7Ph4u) | All | merFISH | single-cell | 5488 | 158 | 81 | 103 | 53 | 56 | 57 |
| Mouse Hypothalamus (layer 110) [[10]](https://paperpile.com/c/xwk5cU/7Ph4u) | All | merFISH | single-cell | 5070 | 156 | 78 | 108 | 53 | 53 | 58 |
| Mouse Hypothalamus (layer -90) [[10]](https://paperpile.com/c/xwk5cU/7Ph4u) | All | merFISH | single-cell | 5557 | 158 | 78 | 98 | 51 | 54 | 55 |
| Mouse Hypothalamus (layer 10) [[10]](https://paperpile.com/c/xwk5cU/7Ph4u) | All | merFISH | single-cell | 5338 | 157 | 75 | 110 | 54 | 56 | 59 |
| Mouse Hypothalamus (layer -290) [[10]](https://paperpile.com/c/xwk5cU/7Ph4u) | All | merFISH | single-cell | 5517 | 156 | 80 | 85 | 46 | 45 | 48 |
| Mouse Hypothalamus (layer 260) [[10]](https://paperpile.com/c/xwk5cU/7Ph4u) | All | merFISH | single-cell | 4832 | 153 | 73 | 87 | 43 | 43 | 44 |
| Mouse Hypothalamus (layer 210) [[10]](https://paperpile.com/c/xwk5cU/7Ph4u) | All | merFISH | single-cell | 4787 | 153 | 76 | 101 | 48 | 52 | 53 |
| Mouse Hypothalamus (layer 160) [[10]](https://paperpile.com/c/xwk5cU/7Ph4u) | All | merFISH | single-cell | 5169 | 155 | 83 | 108 | 56 | 55 | 60 |
